# Malaria control dynamics may explain inconsistent outcomes from bednet trials: a modeling study

**DOI:** 10.1101/435438

**Authors:** James Orsborne, Thomas Walker, Laith Yakob

## Abstract

**BACKGROUND:** Long-lasting insecticidal bednets have unparalleled efficacy in reducing malaria burden. However, insecticidal resistance and bednet avoidance behaviors among the mosquito vectors are now widespread.

**METHODS:** Reviewing the relevant field and semi-field studies highlights the ubiquity of zoophagic and spatiotemporal (biting outdoors or at different times of day) plasticity among vectors in response to bednet deployment. Transmission models coupled with the population genetics of vectors are developed to assess the impact on malaria control caused by insecticide resistance and the avoidance behaviors of mosquitoes.

**RESULTS:** Interactions between physiological resistance and behavioral resilience among mosquito vectors can significantly impact malaria control efforts both in the short- and long-term. The possibility of misleading observations from injudiciously timed assessments of malaria control programs is demonstrated through simulation.

**CONCLUSIONS:** Currently, there are no guidelines to inform when during a bednet trial its effectiveness should be measured. The importance of this oversight is described in the context of recent randomized controlled bednet trials.

## Background

Bednets impregnated with long-lasting insecticide have considerably reduced malaria burden since their scale-up in 2000 ^1^. However, these successes are not ubiquitous. Several bednet programs have described disappointing results ranging from very short-lived health benefits ^2^, to limited reductions in malaria cases ^3–5^, through to abject failure ^6^. More recently, a series of trials investigated the potential additional benefit of complementing bednets with indoor residual spray ^7–9^. Reports again have described contradictory findings, whereby this integrated vector management strategy resulted in anything from synergism to antagonism, relative to bednets alone. These important inconsistencies have been attributed to differences in coverage levels, health systems and vector ecology ^10^. However, post hoc explanations have thus far been anecdotal, lacking the necessary framework for rigorous assessment.

There are many aspects of vector ecology that would be expected to impact efficacy of insecticide-based control tools used exclusively indoors ^11^. Chief among them are the development of resistance to insecticides and the biting behavior of local vectors. Until 2017, the pyrethroids were the only class of insecticides with World Health Organization (WHO) approval for use on bednets ^12^ and their extensive deployment has inevitably led to the emergence and rapid spread of pyrethroid resistance across Africa ^13^. Numerous mosquito populations across Africa and Asia have also evolved resistance to insecticides used in alternative vector control methods such as indoor residual spray ^14^. Currently unclear is whether behavioral changes observed in the major malaria vectors also have a genetic basis. These changes include higher rates of outdoor biting (exophagy) and feeding times altered to when people are not sleeping under bednets ^15, 16^, hereafter these behaviours are grouped and referred to as ‘spatiotemporal plasticity’. They also include increased rates of feeding on animals (zoophagy) in response to high coverage with indoor control tools ^17^.

To supplement the WHO Global Plan for Insecticide Resistance Management, new guidelines for countries to develop their own resistance management plans will soon be published ^18^. The current study describes the development of a mechanistic framework for assessing epidemiological impact of insecticide resistance and adaptive malaria vector behavior. Results will be discussed in the context of recent bednet effectiveness trials to inform developments in country-level resistance management strategy.

## Results

Evidence for bednets impacting mosquito behavior was collated. Synthesising all published studies of the key African malaria vectors (*Anopheles gambiae* and *An*. *funestus*) demonstrated a marked reduction in the Human Blood Index (‘HBI’ is the proportion of blood meals of human origin ^19^) following a bednet program (Figure 1A). Formal statistical assessment of this increased zoophagy was precluded by the fact that the limited number of studies reporting HBI before and after bednets differed so substantially in how long after distribution they conducted their follow-up ^20–27^.

**Figure 1.**
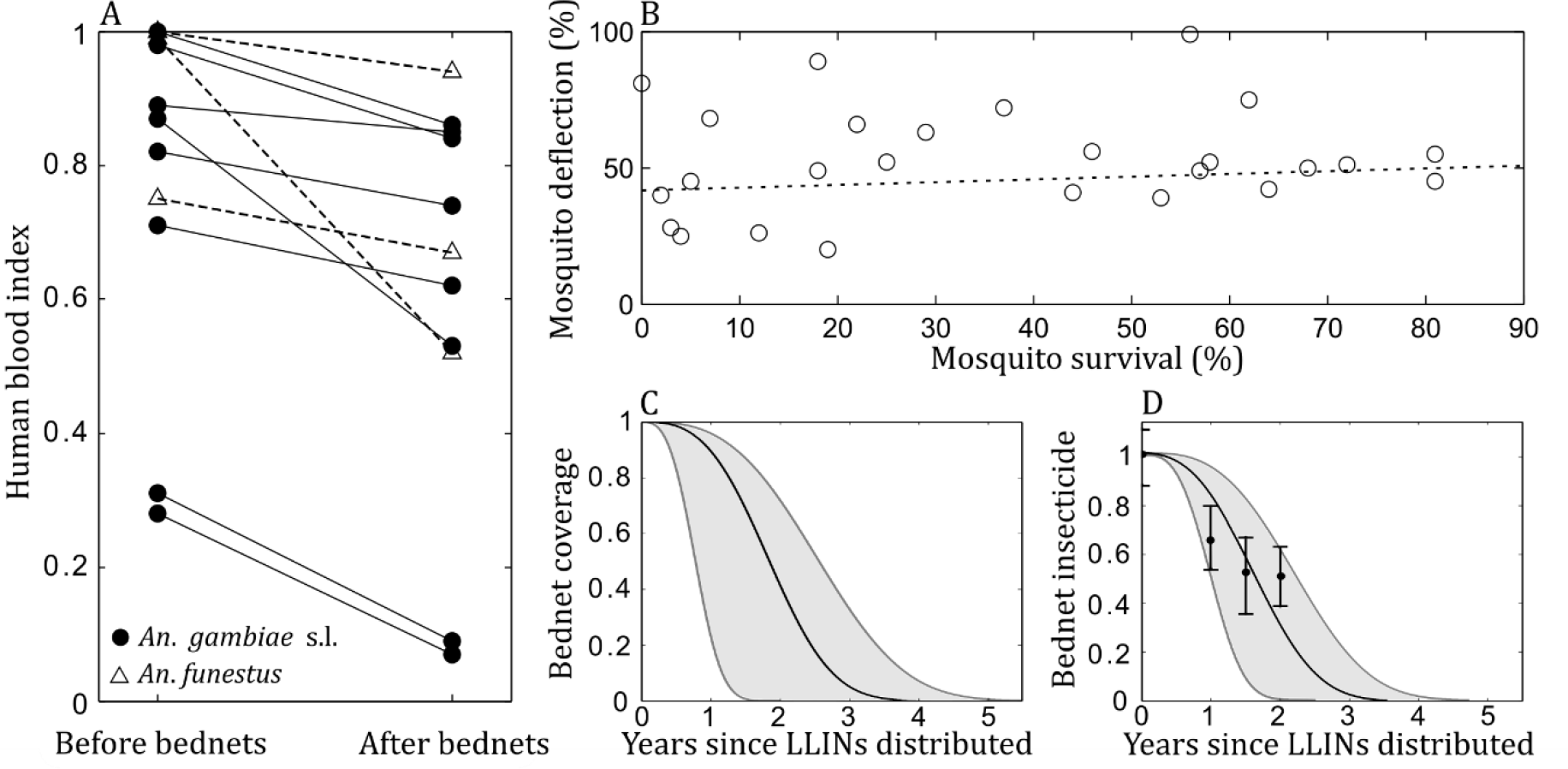
Empirical studies used to inform entomological components of the model. A) Synthesis of studies measuring the human blood index before and after bednet distributions. B) The association between mosquitoes seeking bloodmeals away from dwellings using bednets and their surviving insecticides, as recorded in hut trials (re-analysis of data collated in ^29^). No statistically significant association was found between spatiotemporal plasticity and sensitivity to insecticides: t-value = 0.65, df = 25, p = 0.523. C) The decline in median (and range) of effective coverage with bednets over time as reported in African studies (data reviewed previously ^30^). D) The decline in insecticidal concentration of bednets over time as reported in a longitudinal study in Kenya ^31^.

Evidence for possible interactions between physiological resistance and behavioral adaptations among malaria vectors was also assembled. Largely because of the operational difficulties involved in measuring mosquito behavior ^28^, field studies describing both these factors are scant. However, hut trials of bednet efficacy report insecticide resistance frequency and bednet avoidance behavior as standard. Re-analysis of hut trial data collated in a recent systematic review ^29^ demonstrated no evidence for any association between insecticide sensitivity and propensity of mosquitoes to avoid entry of, or rapidly exit, huts where new bednets were in use (t-value = 0.65, df = 25, p = 0.523; see Fig 1B).

Bednets wane in their effectiveness over time as a function of three components: usage rates decline, the netting material becomes degraded and the potency of the chemical insecticide fades. Respectively, Fig 1C and 1D depict bednet effectiveness curves as derived from usage and netting material integrity data reviewed by Bhatt et al. ^30^ and recently published longitudinal insecticidal concentration data from western Kenya ^31^. These data were used to inform the structure and parameterization of a mathematical model (see *Materials and Methods*). The model was designed to evaluate the dynamic impact of vector behavior both on the spread of insecticide resistance and on malaria control, over the lifetime of a bednet across several rounds of distribution.

Bednet avoidance (‘spatiotemporal plasticity’) reduces selection pressure and thereby delays the spread of physiological insecticide resistance (Fig 2A). However, accounting for this entomological behavior not only reduces epidemiological impact of bednets but also delays impact, with the greatest reduction in parasite rate trailing by up to 7 months in simulations (compare the delayed decline in parasite rate for high-versus low-level spatiotemporal plasticity in Fig 2A). The inclusion of zoophagic plasticity further exaggerates temporal differences in infection control, with the time until achieving maximum control varying by over a year (Fig 2B). Importantly, even in the presence of a high level of insecticide resistance, and regardless of vector behaviors, parasite rates were significantly reduced following bednet distributions.

**Figure 2.**
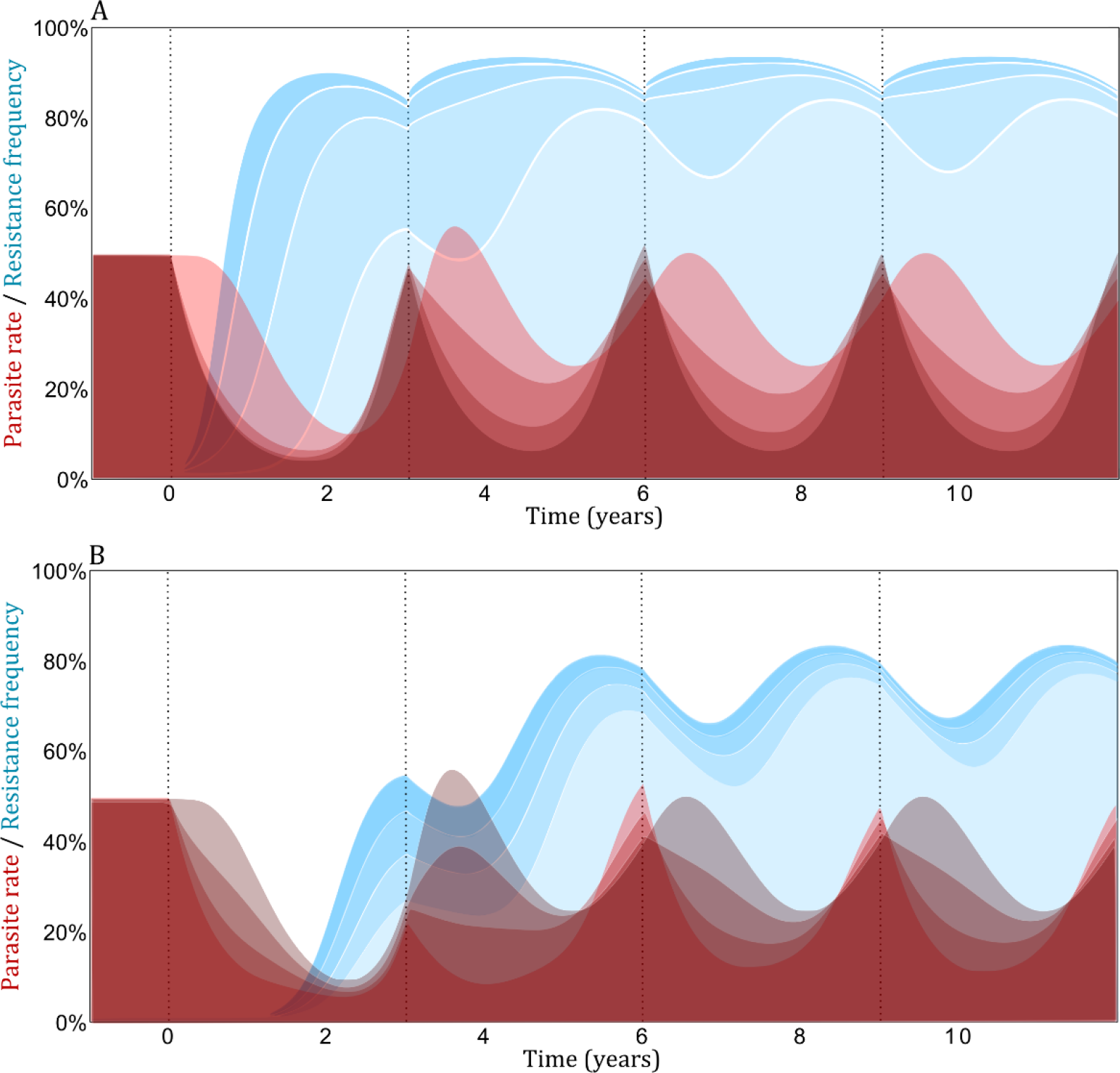
Parasite rate and insecticide resistance frequency during a bednet program with nets redistributed every 3 years (dashed vertical lines). Four different scenarios are presented whereby the brighter reds and lighter blues are associated with greater A) spatiotemporal plasticity (*χ* = 0, 0.33, 0.66, 1; zoophagic plasticity set to 0); or B) zoophagic plasticity (*ζ* = 0, 0.33, 0.66, 1; exophagic plasticity set to 1). Simulations were initiated with low resistance frequency (1%) and an endemically stable 50% parasite rate which was achieved by adjusting parameter *m* (see *Materials and Methods*). Results are illustrative and presented for baseline parameter estimates (see Table 1); uncertainty analysis in these point estimates is conducted later (see *Materials and Methods*).

The combined impact of spatiotemporal and zoophagic plasticity produced outcomes that were mostly intuitive: infection control is more compromised by vectors that avoid bednets and yet remain highly anthropophagic; and, resistance spreads less rapidly when mosquitoes exhibit spatiotemporal and/or zoophagic plasticity (Fig 3). The difference that either behavior can make in terms of disease control and resistance spread is contingent on the intensity of resistance. ‘Resistance intensity’ refers to the strength of resistance i.e. by how much the additional mortality incurred by insecticides is attenuated in ‘resistant’ versus ‘sensitive’ mosquitoes ^32^. When resistance intensity is low, a special case arises whereby bednet efficacy can actually be reduced by a more zoophagic vector when it exhibits only limited spatiotemporal plasticity (see bottom half of Fig 3E). Here, the reduced mosquito mortality due to bites being redistributed away from humans (and, thereby bednets) more than offsets the reduced contact rate with humans, and there is a net increase in the force of infection.

**Figure 3.**
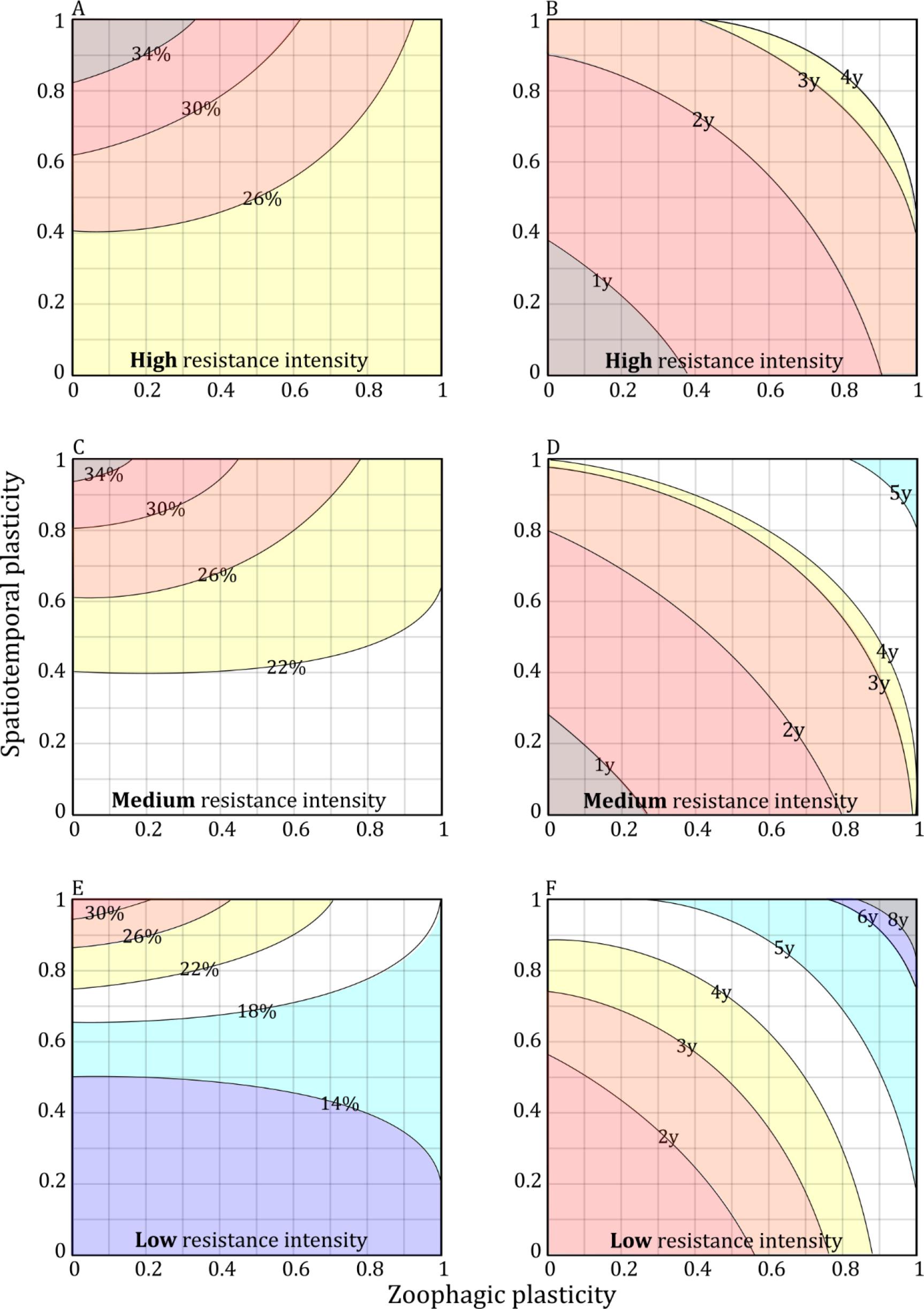
Impact of spatiotemporal and zoophagic plasticity on malaria infection prevalence (averaged over the fourth bednet distribution cycle; left column: 3A, C and E) and on the number of years (y) following bednet distribution before insecticide resistance reaches 50% frequency in the local vector population (right column: 3B, D and F). The resistance intensity is varied with row (3A&D: 10-fold reduced mortality, 3C&D: 5-fold, 3E&F: 2.5-fold). Other parameterization equivalent to Fig 2.

How these entomological behaviors impact epidemiological effectiveness estimates in bednet trials is illustrated in Figure 4. Recent trials in Sudan ^33^, Kenya ^34^, Tanzania ^9^, The Gambia ^7^ and Benin ^8^ differed in the time after bednet distribution at which malaria control impact was assessed (range: 9-18 months). Projections of the mathematical model parameterized with the best data available on bednet longevity demonstrate that this inconsistent time point for measuring control effectiveness invalidates comparisons between trials (Fig 4). For example, if after bednet distribution the local mosquito population exhibited spatiotemporal plasticity but remained highly anthropophagic, the time point at which malaria control was assessed in the Sudanese, Kenyan and Tanzanian studies would have yielded estimates that were *worse* than no control. However, had effectiveness not been measured until later in the trial (as was conducted in The Gambia and in Benin), it would have demonstrated a health benefit of the interventions.

**Figure 4.**
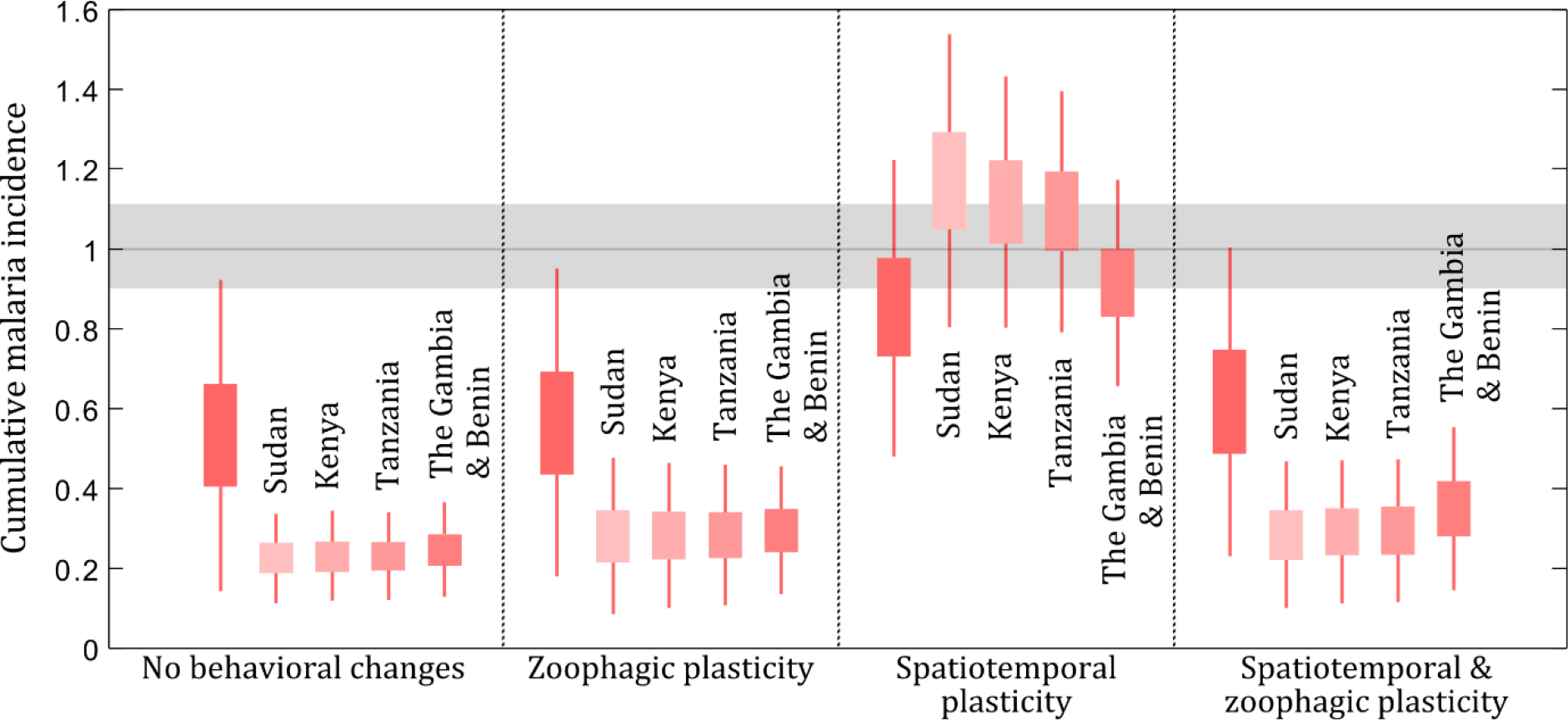
Timing can impact the assessment of bednet effectiveness. Recent bednet trials in Sudan, Kenya, Tanzania, The Gambia and Benin differed in the time after distribution at which malaria control impact was assessed. The cumulative malaria incidence up until time points corresponding with the trial assessments are plotted. For comparison, the cumulative malaria incidence at the end of a three-year bednet cycle is also shown (dark red). The horizontal grey line and band respectively indicate the no-control median and the first and third quartiles of simulations from 1000 random parameter sets. Similarly for the boxplots which also have whiskers representing the interquartile range. Four separate entomological scenarios are depicted in X-axis labels (left to right: *χ* = 0 and *ζ* = 0; *χ* = 0 and *ζ* = 1; *χ* = 1 and *ζ* = 0; *χ* = 1 and ζ = 1).

## Discussion

Reports of bednets becoming less effective in controlling malaria are emerging and no alternative tool of equivalent potential is anticipated in the near future. It is essential to make best use of bednets to prolong their effectiveness; but to achieve this, a better understanding of their ecological impact on mosquito vectors is needed.

Models have previously explored the impact of behavioral plasticity ^35–38^ and population genetics of insecticide resistance ^39–42^ among mosquitoes exposed to bednets. Some reassurance is gained from the current analysis which indicates that reductions in malaria are anticipated even in settings with high plasticity/resistance - a result that is not only corroborated by previous modeling studies but also the majority of empirical studies ^43^. However, while resistance and behavioral adaptations do not appear to completely negate malaria control by bednets, this analysis suggests that these phenomena can significantly attenuate control effectiveness. For a control tool that is estimated to save in the order of half a million lives per year ^1^, diminished returns may signal an imminent global health disaster.

Understanding how behavior and physiological resistance interact is important because spatiotemporal and zoophagic plasticity reduce the selection pressure for resistance development. This means that although malaria control is diminished by an increased tendency for mosquitoes to bite outdoors or at different times of day, the level of control that is achieved is expected to be maintained for longer because resistance development is delayed as a result. Longitudinal data on the HBI and from human-landing catches performed alongside epidemiological assessments are needed to inform projections of bednet effectiveness both in the short- and long-term.

To monitor resistance, WHO guidelines recommend exposing insects to a discriminating dose of insecticide for a set period of time and recording the percentage mortality. While indicative of general spatial and temporal trends, findings from these bioassays are not easily converted into metrics useful to field applications. For example, how does the percentage of mosquitoes dying after 60 minutes of forced exposure to 0.75% permethrin in a test tube inform the bednet-associated mortality of mosquitoes in the field? New bioassay methods to quantify resistance intensity have been developed to improve this translation ^32^, but are so labor-intensive as to preclude their use in large-scale trials assessing the public health impact of insecticide resistance ^44^. Churcher et al. ^29^ sought associations between resistance as measured by mosquito bioassays and results from hut trials. Once validated, these associations will require a second, more complex, conversion from semi-field conditions to real-world settings e.g. to account for opportunities for vectors to source blood meals outdoors and from alternative host species.

Results presented here indicate that the interaction between behavioral plasticity and physiological resistance is moderated by resistance intensity. Saul ^36^ described the potential for zooprophylaxis to switch into zoopotentiation if the availability of alternative blood meals increases mosquito survival more than counters the impact of diverting feeds. This scenario is also shown in the current model’s projections but here zoopotentiation only resulted when resistance intensity was low (insecticide associated mortality in resistant mosquitoes is only 2.5 fold less than for sensitive mosquitoes). Should data become available for behavioral adaptations corresponding with data on resistance dynamics and intensity, the current framework will be able to assess the risk of zoopotentiation.

Disentangling these interactions will become even more important in the context of systemic insecticides: drugs that render host blood toxic to haematophagous ectoparasites such as mosquitoes. Recently, there has been growing interest in the use of systemic insecticides such as ivermectin ^45–47^ and fipronil ^48 49^ to target both livestock- and human-biting mosquitoes; and there have been developments in models to inform strategic use of these tools as part of the integrated vector management of malaria vectors ^38 50^. Future work utilising the current framework is needed to assess malaria-control impact of bednets used in conjunction with these and other tools. Recent trials of malaria vector control have been dominated by assessments for the combined impact of IRS with LLINs and it will be critical to ascertain how the conclusions of this study are altered for an integrated vector management programme.

There are several additional aspects of the current framework that can be developed further. As with all models, a trade-off exists whereby the complexity of additional realism compromises transparency. Despite there being several mechanisms of resistance in malaria vectors ^51^, only a single trait acting in isolation is considered here. Metabolic resistance to pyrethroids is generally considered the greatest threat to operational success ^52^ and this was the mechanism focused on in the current study. Like other population genetics studies of insecticide resistance among malaria vectors, resistance is assumed to be determined by a single-locus allele spread through random mating ^53^. Recently, Levick et al. developed a two-locus model of insecticide resistance evolution for malaria vectors ^54^ demonstrating the considerable increase in model complexity required by this addition. As greater understanding is gained of the numerous mechanisms and how they potentially interact ^55 56^, or if strong evidence arises for a genetic basis of behavioral plasticity among the main malaria vectors, a more comprehensive genetic component may become an important future adaptation, and even the two-locus model may need further extension.

An additional simplification made to improve model transparency involves the processes governing infection. These are driven by a Susceptible-Infected-Recovered-Asymptomatically Infected system (see *Materials and Methods*) when, in reality, the immunology of malaria infection is extremely complicated and incompletely understood. Projections should, as ever, be interpreted with caution. Models such as the one presented here are useful at identifying important absences in data and for contributing towards strategy-level recommendations. For projections to become more operationally suitable, location-specific epidemiology and ecology must be taken into account and a more complex model will be justified.

## Conclusions

For all recent bednet trials, the primary endpoint of clinical malaria cases was measured at a follow-up period that was as much determined by the need to report findings within fleeting project lifetimes as it was by the epidemiology of infection post-control. This absence of consistently collected, coupled entomological-epidemiological data spanning the recommended duration of a bednet distribution round represents much more than a limitation in projection fitting; it highlights large disparities between trials of bednet effectiveness. The timescale across which effectiveness should be measured and optimised has not been formalized. Here, through simulation it is demonstrated that even in the highly unrealistic situation that the recent bednet trials were conducted identically in epidemiologically and entomologically indistinguishable locations, the different time points at which they were assessed would potentially be sufficient to generate qualitative differences in their recorded impact.

Following promising results using a bednet impregnated with Chlorfenapyr ^57^, it is likely that this will become the first new insecticidal class to receive WHO approval for use in bednets in decades. Strategic targeting of this precious new tool is paramount. Although no cross-resistance to this class of active ingredient is anticipated, this new generation of bednets will be at the mercy of extant vector behavioral adaptations. Findings from the current study are hoped to help inform more judicious assessment of its effectiveness in controlling malaria.

## Methods

### Systematic review

PRISMA guidelines were followed for the systematic review. The key word search, inclusion and exclusion criteria agreed by authors before the systematic search was performed. The Ovid database was used to search available MEDLINE and EMBASS literature from inception to February 2018. All books were excluded from all searches and only articles written in English were included. Results were collated and managed using Mendeley desktop reference manager.

Search strategy: Human blood index **OR** HBI **OR** host preference **OR** trophic preference **OR** blood meal preference **OR** blood host preference **OR** blood meal **OR** blood meal analysis **OR** blood-meal analysis **OR** blood meal source **OR** host blood **OR** host blood meal **OR** blood meal identification [multiple posting= MeSH subject heading word, abstract, title, original title, text word (title, abstract), key word heading, name of substance, key word heading word, protocol supplementary concept word, synonym]

### AND

Anopheles **OR** Anopheles arabiensis **OR** Anopheles gambiae **OR** Anopheles funestus [multiple posting= MeSH subject heading word, abstract, title, original title, text word (title, abstract), key word heading, name of substance, key word heading word, protocol supplementary concept word, synonym]

Inclusion criteria were: studies which used blood meal analysis (PCR,ELISA or precipitin tests) to report the HBI; HBI reported both before and after bednet distributions; studies performed in sub-Saharan Africa; reporting HBI for *Anopheles gambiae* or *Anopheles funestus* complex. Exclusion criteria were: semi field studies; studies using baited traps or choice experiments to investigate host preference; studies not reporting numbers of mosquitoes caught; studies reporting HBI for fewer than 50 blood-fed mosquitoes. Duplicates were removed and abstracts for all publications retrieved were reviewed for relevance. Full-text reviews were then conducted on all articles. If the inclusion criteria were satisfied the paired estimates for reported Human Blood Index (HBI) were retrieved.

### Mosquito population dynamics

Mosquito population dynamics are described using a time-delay difference equation model, adapted from ^58^. The model explicitly tracks the number of adults (*A*) over time, *t*, while accounting for density dependent survival of the larval mosquito stages:

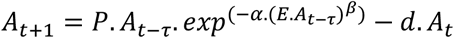

Parameter *P* denotes the per capita reproduction rate corrected for density independent mortality during pre-adult stages. *E* is the per capita daily egg production rate and the adult mortality rate is denoted by *d*. The time-delay component accounts for the generational time (*τ*) of *Anopheles gambiae* (*A*_*t-τ*_ refers to the adult mosquito population *τ* days ago). *α* and *β* respectively govern the carrying capacity of mosquito larvae and the intensity of density dependence. The impact that parameterization of this equation has on stability is explored fully elsewhere ^59^. Briefly, if *β*.ln(*P*/*d*)>1 dynamics are unstable, yielding monotonically dampening cycles, tending towards stable limit cycles and ultimately chaotic dynamics for very high values. Mosquito populations in the field are very unlikely to exhibit chaotic dynamics (this being an extremely rare trait of any natural insect population) and typically tend towards the stability boundary condition ^59–61^. Scenarios modelled here thereby assume parameterization that simulates a stable mosquito population (*β*.ln(*P*/*d*)≈1). However, a range of dynamic behaviors are tested, allowing for parameterization to encompass stable and unstable mosquito population dynamics (see *Uncertainty analysis*).

### Population genetics of insecticide resistance

Typically, models of the spread of resistance assume genotype frequencies follow standard replicator dynamics ^62^. However, because current focus is on the temporal dynamics of resistance and behavioral adaptions to bednets as their efficacy wanes (see *Incorporating vector control* section), each single-locus resistance genotype (*ss*, *sr* and *rr* are abbreviated to superscript *i* below for brevity) was tracked explicitly at each daily time-step for both male (*M*) and female (*F*) mosquitoes:

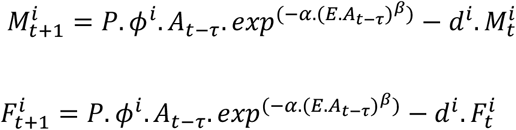

Whereby,

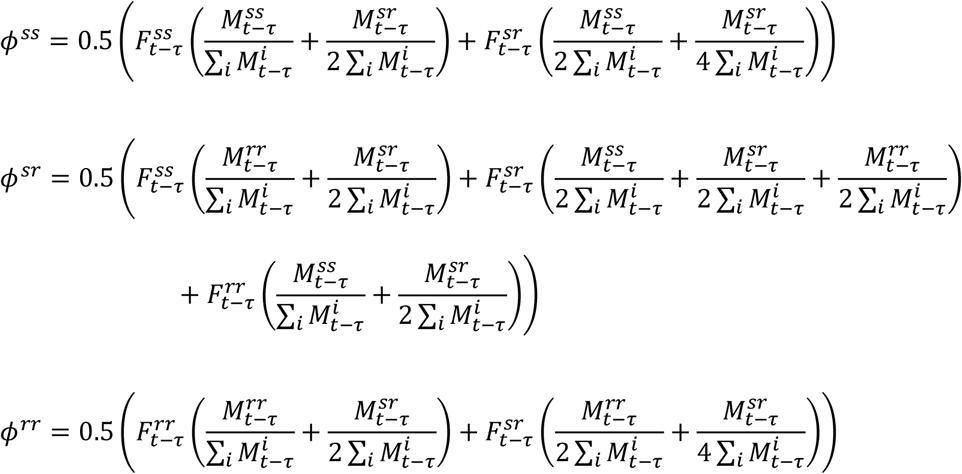

The resistance phenotype was assumed dominant to present cautious estimates of the impact from attenuated bednet efficacy. Fitness costs were conservatively assumed to impact only homozygous resistant mosquitoes, and modeled by incurring an increased mortality rate (i.e. *d*^*rr*^ > *d*^*sr*^ & *d*^*rr*^).

### Malaria infection transmission

Malaria infection dynamics were simulated using the following system of difference equations:

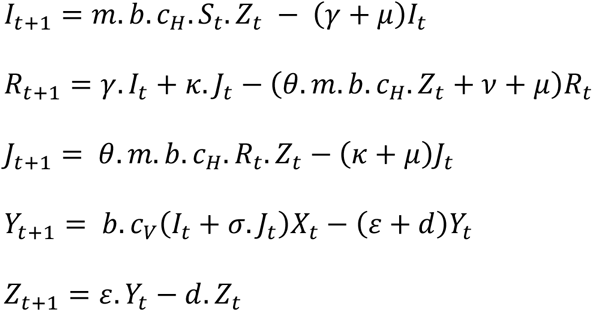

People are susceptible (*S*), infected (symptomatic, *I*), recovered (*R*) or asymptomatically infected (*J*); and a stable population is assumed: *S*=1-(*I*+*R*+*J*). Infections were split between symptomatic and asymptomatic infections because epidemiological endpoints may differ between trials whereby effectiveness may be measured in terms of reduced incidence (i.e. *I* only) or reduced parasite rate (i.e. *I* plus *J*). It is assumed that an asymptomatic infection can only arise in an individual that has recently been infected already.

Following convention, *m* represents the ratio of mosquitoes to humans; *b* represents the bite rate and *c* is the transmission coefficient (subscript ‘*H*’ denotes transmission from vector to host; subscript ‘*V*’ denotes transmission from host to vector). The human mortality rate is denoted *μ* (malaria associated mortality was not included). Respectively, *γ* and *κ* are the rates of recovery from symptomatic and asymptomatic infection. The rate of loss of immunity is *ν*. Reduced susceptibility of recovered individuals to secondary infection is accounted for by *θ*.

Mosquitoes are susceptible (*X*), infected but not yet infectious (*Y*) or infectious (*Z*). In the absence of control, a stable mosquito population is assumed: *X*=1-(*Y*+*Z*). *σ* allows for a different level of parasite transmissibility from asymptomatic individuals to mosquitoes (relative to symptomatic individuals); and *∊* is the reciprocal of the extrinsic incubation period for the parasite. Parameter definitions and sources for their values are described in Table 1.

**Table 1.**
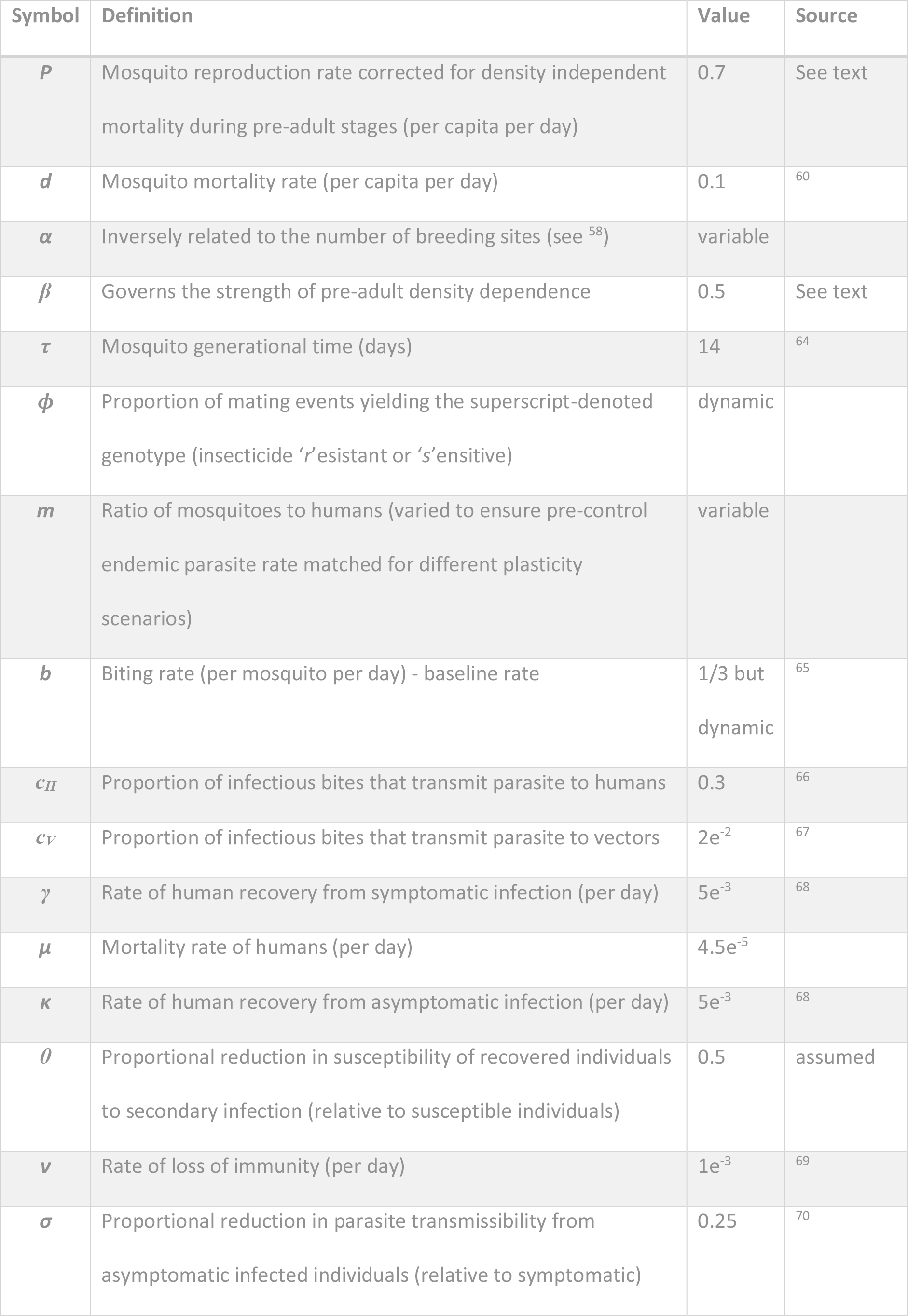
Model parameter definitions, values and empirical sources.

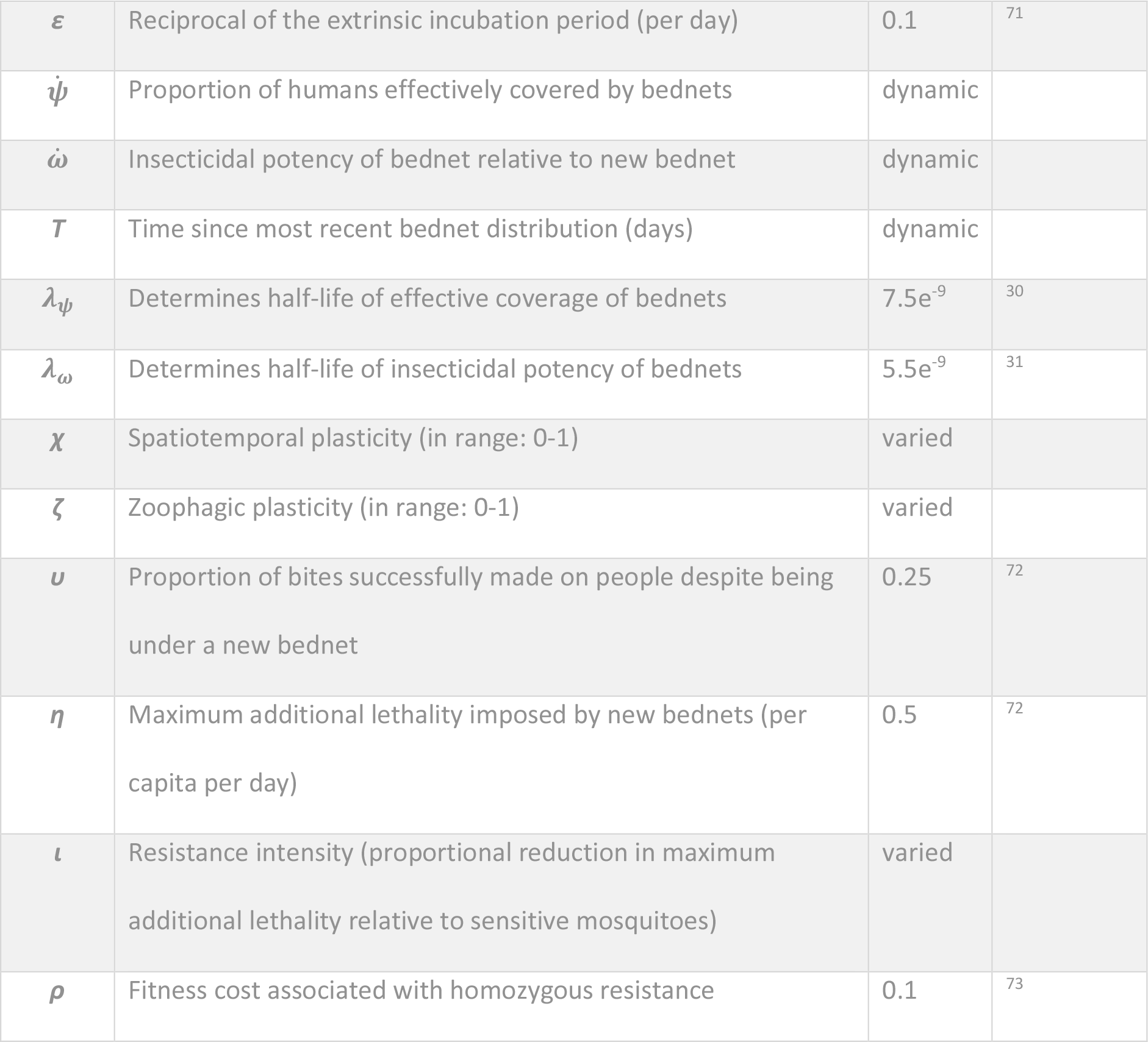

### Incorporating vector control

Bednets reduce the bite rate on humans and kill mosquitoes that come into contact with the impregnated insecticide. Both of these effects wane over time as the net accumulates holes and the insecticide loses potency. Yakob et al. ^63^ developed methods to account for decayed bednet efficacy making use of a squared exponential function that better resembles the initially slow, but accelerating, decline in efficacy over time that is reported empirically ^30^. Here, this is taken further, allowing the decline in usage rates reported over time coupled with the physical degradation of the nets to inform effective coverage 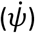:

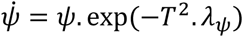

Where, *ψ* is maximum effective coverage of new bednets immediately following their distribution; *T* is time (in days) since the most recent bednet distribution; and *λ*_*ψ*_ informs the efficacy half-life. An equivalent function is used to describe the waning in insecticidal content of the bednet over time 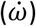. Universal distributions of new bednets are simulated to occur every 3 years in line with WHO guidelines. For simplicity, any additional control achieved by older nets surviving between distributions is ignored. Again, this reflects a conservative scenario; in the future, this simplification could be interrogated by behavioral data on longer term bednet use.

The impact that bednets have on the bite rates and mosquito mortality is not only affected by their effective coverage and insecticidal potency but also by spatiotemporal and zoophagic plasticity in the vector. Brand new bednets act as powerful physical and chemical barriers and so the pressure on the mosquitoes to bite outdoors, at a different time of day or a non-human host is at its greatest immediately following bednet distribution. These pressures wane over time along with net integrity/usage and insecticidal potency. Hence, it is assumed that the effective human-biting rate 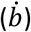 is impacted less as the bednets age:

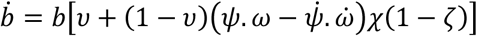

Where *χ* and *ζ* are the level of spatiotemporal and zoophagic plasticity respectively (both in the range 0-1). In this way, mosquitoes that are less spatiotemporally plastic waste more time trying to secure bloodmeals from humans resting under bednets and consequently make fewer bites over their lifetime. However, the magnitude of this effect is attenuated by the proportion of bites successfully made on people despite being under a new bednet 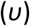.

Similarly, 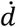 denotes mosquito mortality incorporating the effects of vector control and is included thusly:

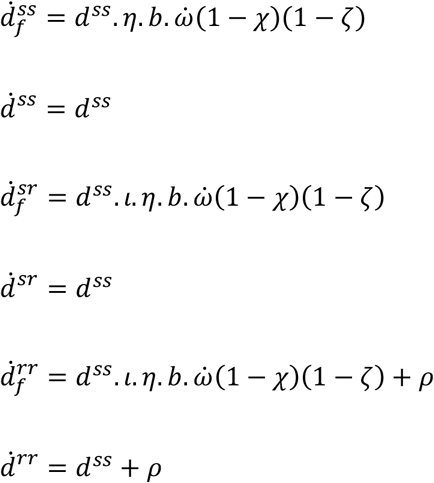

Where *η* is the maximum additional lethality imposed by new bednets, and this is attenuated for both heterozygous and homozygous resistant mosquitoes by factor *ι* which is the resistance intensity. This assumes complete dominance of the resistance allele, simulating the most conservative scenario. Should evidence arise for incomplete dominance of resistance alleles in major malaria vectors, this can easily be incorporated into the model by multiplying the additional mortality for heterozygous females by the level of dominance ^62^. Parameter *ρ* is the fitness cost associated with resistance and, conservatively, is only assumed to penalize homozygous insects.

### Uncertainty analysis

Parameter uncertainty was taken into account by allowing all input parameters to vary around the point estimates found in the empirical literature (Table 1). 1000 parameter sets were generated in which each input parameter varied randomly ±50% within a uniform distribution (i.e. a distribution that conservatively does not assume skewed central tendency around the point estimate). These random parameter sets were used to test the robustness of results comparing the different timings at which different bednet trials conducted effectiveness assessments.

## Declarations

### Ethics approval and consent to participate

N/A.

### Consent for publication

N/A.

### Availability of data and material

The current study involves the analysis of previously published data only.

### Authors’ contributions

LY conceived the study and constructed the mathematical model; JO, TW, LY contributed towards the systematic review, results interpretation and manuscript drafting.

## Acknowledgements

N/A

